# The type 2 asthma mediator IL-13 inhibits SARS-CoV-2 infection of bronchial epithelium

**DOI:** 10.1101/2021.02.25.432762

**Authors:** Luke R. Bonser, Walter L. Eckalbar, Lauren Rodriguez, Jiangshan Shen, Kyung Duk Koh, Lorna T. Zlock, Stephanie Christenson, Prescott G. Woodruff, Walter E. Finkbeiner, David J. Erle

**Author notes:** **Corresponding Author**: David J. Erle, MD, CVRI - MC: 3118, 555 Mission Bay Blvd South, PO Box 589001, San Francisco, CA 94158-9001. These authors contributed equally (Luke R. Bonser, Walter L. Eckalbar). **Contributions**: LRB, WLE, LR, and DJE contributed to study conception and design. LRB, KDK, LTZ, and LR performed experiments. LRB, WLE, JS, KDK, SC, PGW, and DJE analyzed and interpreted data. WEF and DJE supervised study execution. LRB and DJE drafted the manuscript. All authors reviewed the manuscript. **Support**: NIH awards U19 AI 077439 and R35 HL145235 (to DJE) and R00 HL135403 (to WLE), University of California, San Francisco Program for Breakthrough Biomedical Research (PBBR; to LRB).

## Abstract

**Rationale:** Asthma is associated with chronic changes in the airway epithelium, a key target of SARS-CoV-2. Many epithelial changes are driven by the type 2 cytokine IL-13, but the effects of IL-13 on SARS-CoV-2 infection are unknown.

**Objectives:** We sought to discover how IL-13 and other cytokines affect expression of genes encoding SARS-CoV-2-associated host proteins in human bronchial epithelial cells (HBECs) and determine whether IL-13 stimulation alters susceptibility to SARS-CoV-2 infection.

**Methods:** We used bulk and single cell RNA-seq to identify cytokine-induced changes in SARS-CoV-2-associated gene expression in HBECs. We related these to gene expression changes in airway epithelium from individuals with mild-moderate asthma and chronic obstructive pulmonary disease (COPD). We analyzed effects of IL-13 on SARS-CoV-2 infection of HBECs.

**Measurements and Main Results:** Transcripts encoding 332 of 342 (97%) SARS-CoV-2-associated proteins were detected in HBECs (≥1 RPM in 50% samples). 41 (12%) of these mRNAs were regulated by IL-13 (>1.5-fold change, FDR < 0.05). Many IL-13-regulated SARS-CoV-2-associated genes were also altered in type 2 high asthma and COPD. IL-13 pretreatment reduced viral RNA recovered from SARS-CoV-2 infected cells and decreased dsRNA, a marker of viral replication, to below the limit of detection in our assay. Mucus also inhibited viral infection.

**Conclusions:** IL-13 markedly reduces susceptibility of HBECs to SARS-CoV-2 infection through mechanisms that likely differ from those activated by type I interferons. Our findings may help explain reports of relatively low prevalence of asthma in patients diagnosed with COVID-19 and could lead to new strategies for reducing SARS-CoV-2 infection.

## Introduction

One remarkable feature of the COVID-19 pandemic is the wide range of disease severity seen following SARS-CoV-2 infection. Host factors, including age and male sex (1), inborn or acquired disorders of type I interferon-mediated antiviral immunity (2–4), and various pre-existing medical conditions (5) influence the risk of severe disease. There have been concerns that COVID-19 risks would also be increased in persons with asthma, which affects ∼339,000,000 individuals worldwide (6). These concerns arose from experience with other respiratory viruses which trigger asthma exacerbations and can be associated with worse outcomes in individuals with pre-existing asthma (7). However, asthma was underrepresented in early studies of patients with COVID-19 as well as prior studies of the severe acute respiratory syndrome (SARS) (8). Subsequent studies have failed to find consistent evidence of increased risk of COVID-19 diagnosis, hospitalization, or mortality due to asthma (1, 9, 10), and some studies concluded that risk of acquiring and being hospitalized for COVID-19 are lower in those with asthma (10, 11). Factors that might provide protection against COVID-19 in individuals with asthma (12–15) include increased attention to limiting viral exposure, younger age, absence of co-morbidities, use of inhaled corticosteroids, and a variety of biological features of asthma, including chronic airway inflammation, mucus hypersecretion, and altered expression of SARS-CoV-2 receptor (12). However, direct evidence that these features alter SARS-CoV-2 infection has been lacking.

Asthma is associated with changes in the structure and function of the airway epithelium, a critical site for SARS-CoV-2 infection (16, 17). Airway epithelial gene expression changes attributable to the type 2 cytokine IL-13 are seen in approximately half of individuals with asthma (18, 19). IL-13 stimulation of airway epithelial cells decreases expression of *ACE2*, which encodes the SARS-CoV-2 receptor, and increases expression of *TMPRSS2*, which encodes a transmembrane protease that primes the viral spike protein (20–22). Similar changes are seen in airways of individuals with type 2 high asthma (21, 22). IL-13 has also been reported to protect against other RNA viruses, including respiratory syncytial virus (RSV) (23) and rhinovirus (24), that do not rely on ACE2 and TMPRSS2 for entry, indicating that other IL-13-regulated genes can also protect against viral infection. We therefore hypothesized that IL-13 induces changes in expression of airway epithelial cell genes important in SARS-CoV-2 infection in a large subset of individuals with asthma and that IL-13 reduces susceptibility of these cells to SARS-CoV-2 infection.

### Methods

Additional details are provided in the Online Data Supplement.

### Human bronchial epithelial cell (HBEC) culture

Primary HBECs from 13 individuals listed in Table E1 were cultured at air-liquid interface (ALI) as previously described (25, 26). For cytokine stimulation, cultures were stimulated by addition of cytokines (10 ng/ml) to the basolateral medium (IFN-α: 24 hours, IFN-γ: 24 hours, IL-13: 7 days, IL-17: 7 days). The UCSF Committee on Human Research approved these studies.

### RNA-seq

RNA was isolated from cytokine-stimulated HBECs derived from six individuals and bulk RNA-seq was performed as previously described (27, 28). We previously reported other analyses based upon data from unstimulated cells and cells stimulated with individual cytokines (27, 28); data from cells treated with a combination of IL-13 and IFN-α have not been previously reported. For scRNA-seq, single cell suspensions were generated from HBECs from four of the donors used for bulk RNA-seq and analyzed using the 10X Genomics platform.

### Analysis of gene expression in asthma and COPD

Using gene expression data from studies of asthma (27) and COPD (29) we correlated measures of type 2 activity with IL-13-induced SARS-CoV-2-associated genes in HBECs using Pearson’s correlation coefficient and a linear regression model that adjusted for age and sex.

### SARS-CoV-2 infection

SARS-CoV-2 virus (USA-WA1/2020 strain) was provided by Dr. Melanie Ott and propagated in Vero E6 cells. HBECs were cultured in the absence or presence of IL-13 (10 ng/ml). Mucus was allowed to accumulate for 3 d or was removed immediately prior to viral infection by washing the apical surface with a prewarmed solution of 10 mM dithiothreitol (DTT; Thermo) in PBS for 10 minutes (26) and then washing twice with PBS without DTT. Cells were inoculated by adding virus to the apical surface. After 2 h, the apical surface was washed with twice with PBS and cells were returned to the incubator. Cells were harvested 48 h post-infection for analysis of viral RNA (by qRT-PCR) and dsRNA-staining (by immunofluorescence).

## Results

### SARS-CoV-2-associated host genes are highly expressed in HBECs

We compiled a list of 342 SARS-CoV-2-associated genes belonging to one of two gene sets. One set of 11 genes encode proteins implicated in viral entry: *ACE2* (30), *TMPRSS2* (30), the cathepsins *CTSB* and *CTSL, FURIN(PCSK3)*, and the furin-like proteases *PCSK1, PCSK2, and PCSK4*-7 (31). The second set of 332 genes encode host cell proteins shown to interact with high confidence with 26 of the 29 SARS-CoV-2 proteins in HEK293T cells (32). One gene, *PCSK6*, belonged to both sets. Using bulk RNA-seq, we detected 332 of the 342 SARS-CoV-2-associated genes (97% ≥ 1 read per million mapped reads [RPM] in ≥50% of samples) in differentiated HBECs cultured without cytokine (Table E2). SARS-CoV-2-associated genes were substantially overrepresented among genes with high read counts (Fig. 1A).

**Figure 1.**
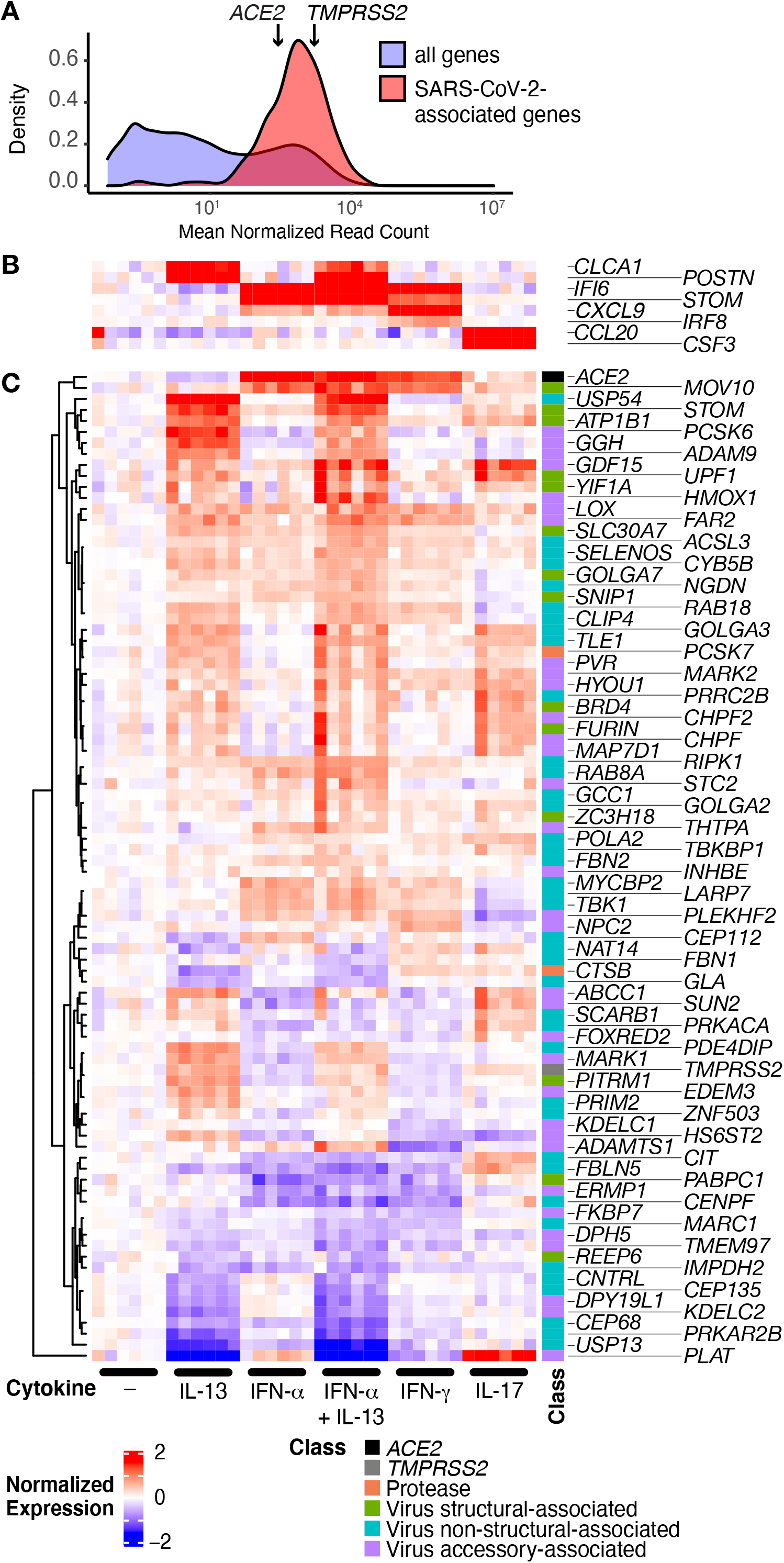
SARS-CoV-2-associated genes are highly expressed in HBECs and many are regulated by cytokines. HBECs from six donors were cultured without cytokine (–), or with IL-13, IFN-α, a combination of IL-13 and IFN-α, IFN-γ, or IL-17 and analyzed by RNA-seq. (**A**) Comparison of read counts between SARS-CoV-2 associated genes, including *ACE2* and *TMPRSS2*, and all detected genes (≥1 read per million mapped reads in ≥50% of samples) in unstimulated HBECs. (**B, C**) Heatmap illustrating canonical cytokine-regulated genes (B), and cytokine regulated SARS-CoV-2-associated genes (C; FDR *q* ≤0.05; absolute fold change ≥ 1.5 for any cytokine).

### IL-13 and other cytokines regulate expression of many SARS-CoV-2-associated genes

We examined the effect of IL-13 stimulation on SARS-CoV-2-associated gene expression. We also analyzed the effects of IFN-α, which plays a central role in defense against SARS-CoV-2 infection (3, 4) and has been shown to inhibit SARS-CoV-2 infection of a human lung epithelial cell line (33), and IFN-γ (27) and IL-17 (34), each of which have been implicated in subsets of individuals with asthma. Each cytokine had the expected effects on expression of known cytokine-responsive genes (Figure 1B). Of the 332 SARS-CoV-2-associated genes detected in HBECs, IL-13 regulated 41 genes, IFN-α regulated 19 genes, the combination of IL-13 and IFN-α regulated 63 genes, IFN-γ regulated 14 genes, and IL-17 regulated 21 genes (false discovery rate [FDR] *q* ≤ 0.05 and absolute fold change ≥ 1.5; Figure 1C; Table E2). *ACE2* was reduced by IL-13 (29% decrease, *q* = 0.003), consistent with a prior report (24). In contrast, *ACE2* was the most highly upregulated gene following stimulation with IFN-α (451% increase, *q* = 3 × 10^−74^) or IFN-γ (185% increase, *q* = 9 × 10^−29^) and was less strongly upregulated following IL-17 stimulation (31% increase, *q* = 0.02). Analysis of *ACE2* splicing confirmed prior reports that interferon stimulation increased expression of the decoy isoform (35), but we found no significant effect of IL-13 on levels of this isoform (mean decoy *ACE2* normalized reads: unstimulated, 62; IFN-α-stimulated: 588, IL-13-stimulated, 61). *TMPRSS2* was increased by IL-13 (61% increase, *q* = 7 × 10^−15^) and IL-17 (22% increase, *q* = 0.005) but modestly decreased by IFN-α (17% decrease, *q* = 0.01) and IFN-γ (16% decrease, *q* = 0.03). No significant correlation between IL-13-and IFN-α-induced changes in the 342 SARS-CoV-2-associated genes was observed, and combined stimulation with both cytokines resulted in additive effects with no evidence of synergy or antagonism (Figure E1). This suggests that these two cytokines affect SARS-CoV-2-associated genes by different and independent mechanisms.

### Expression of many SARS-CoV-2-associated genes differs between airway epithelial cell types

We used single cell RNA sequencing (scRNA-seq) to assess cell type-specific expression of SARS-CoV-2-associated genes in unstimulated HBEC cultures from four individuals (Figure E2). Of the 332 SARS-CoV-2-associated genes detected in HBECs by bulk RNA-seq, 322 (97%) were detected in at least 10 cells. We detected *ACE2* in 1.7% of basal cells, 3.1% of secretory cells, and 1.6% of ciliated cells from unstimulated HBEC cultures. 113 of the 322 SARS-CoV-2 genes detected in our scRNA-seq dataset were differentially expressed between cell types (FDR *q* < 0.05; Table E3; selected genes shown in Fig. 2A). We found similar patterns of epithelial cell subset-specific expression of these SARS-CoV-2 associated genes in a scRNA-seq dataset from human bronchial tissue (36) (Figure E3), confirming that our cell culture model recapitulated cell type-specific gene expression seen *in vivo* and that the expression of host cell proteins that interact with SARS-CoV-2 proteins differs between cell types.

**Figure 2.**
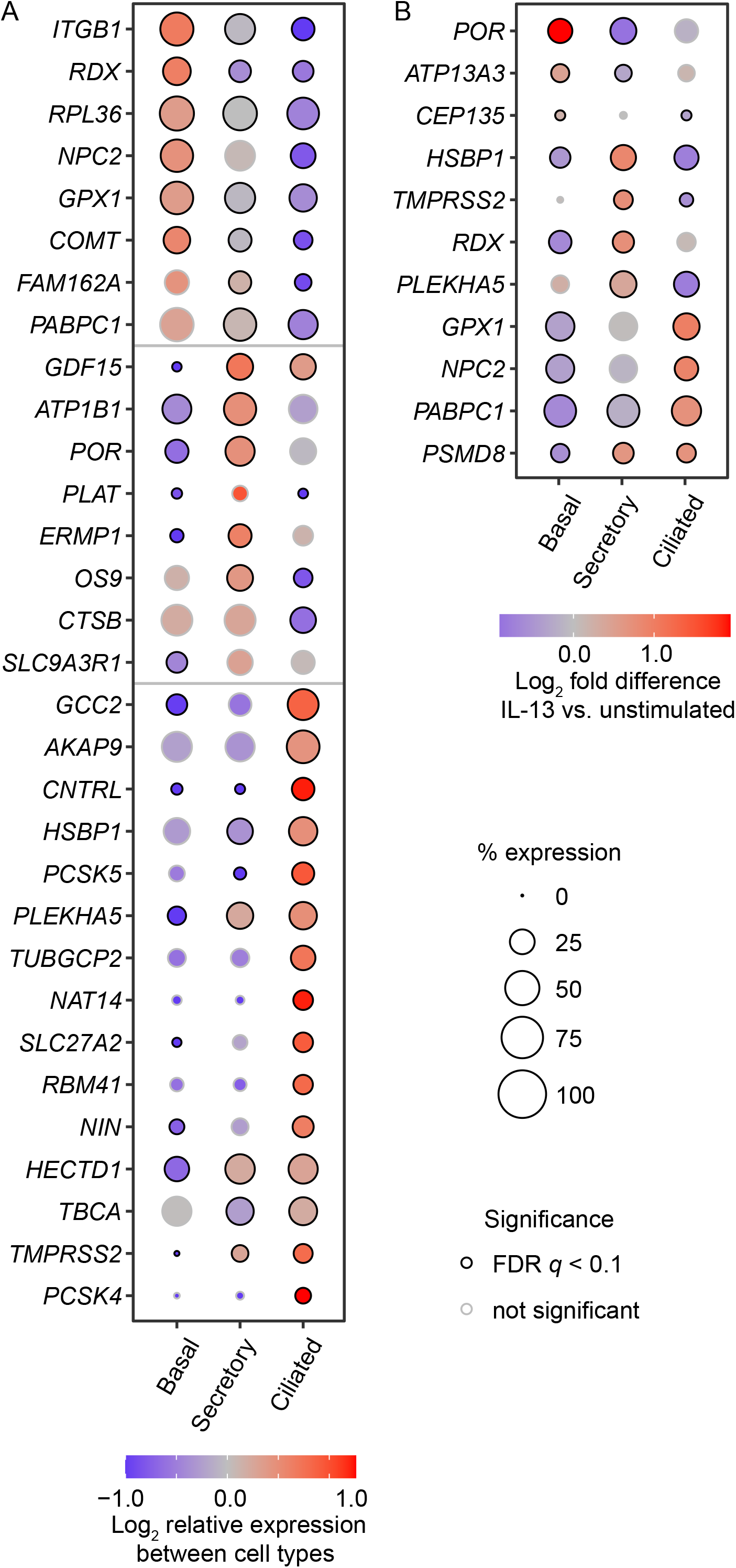
scRNA-seq reveals cell type-specific expression of many SARS-CoV-2-associated genes and cell type-specific effects of IL-13. **(A)** Cell type-specific expression in unstimulated HBECs. Genes were selected from a set of 113 differentially expressed SARS-CoV-2-associated genes listed in Table E3. (**B**) Cell type-specific differences in IL-13 responses. For 11 SARS-CoV-2-associated genes, IL-13 increased expression in at least one cell type and decreased expression in at least one other cell type (FDR *q* < 0.1 for both). Gene expression was determined by aggregating data from all cells from experiments with four donors. Coloring of each dot indicates expression level relative to other cell types (A) or in IL-13-stimulated cells compared with unstimulated cells (B). The size of each dot is proportional to the percentage of cells with at least one read mapped to the gene, and black circles at the perimeter of each dot indicate that expression levels are significantly different (*q* < 0.1) compared to other cell types (A) or in IL-13-stimulated compared to unstimulated cells of the same type (B).

### IL-13 affects expression of some SARS-CoV-2-associated genes in a cell type-specific manner

We explored the effect of cytokine stimulation on SARS-CoV-2 associated gene expression in each cell type. Whereas many IL-13-regulated SARS-CoV-2 associated genes were affected similarly in each cell type, some IL-13 effects were cell type-specific, with 11 cases in which IL-13 had opposite effects (increased in one cell type and decreased in another cell type, FDR *q* < 0.1 for both; Figure 2B and Table E3). Notably, IL-13 upregulated *TMPRSS2* expression in secretory cells, but decreased expression in ciliated cells. Cell type-specific effects of IL-13 could have implications for the outcome of infection in different airway epithelial subsets.

### Type 2 signatures are associated with expression of many SARS-CoV-2-associated genes in individuals with asthma and COPD

We examined whether IL-13-induced SARS-CoV-2-associated genes identified in HBECs in culture were altered in asthma using a transcriptomic profiling dataset derived from endobronchial brush biopsies from individuals with mild to moderate asthma and healthy individuals (Mechanisms of Asthma STudy [MAST]) (27, 37). We used the three gene metric (TGM), an established measure of IL-13-induced airway inflammation in individuals with asthma (18, 19, 38), for this analysis. 24 of 27 SARS-CoV-2-associated genes induced by IL-13 in HBECs were positively correlated with the TGM (Pearson’s *R* > 0); in 13 cases this correlation was statistically significant after adjustment for multiple comparisons (adjusted *p* < 0.05; Figure 3; Table E4). 16 of 27 IL-13-induced genes were significantly associated with the TGM using a linear model that included age and sex (adjusted *p* < 0.05; Table E4).

**Figure 3.**
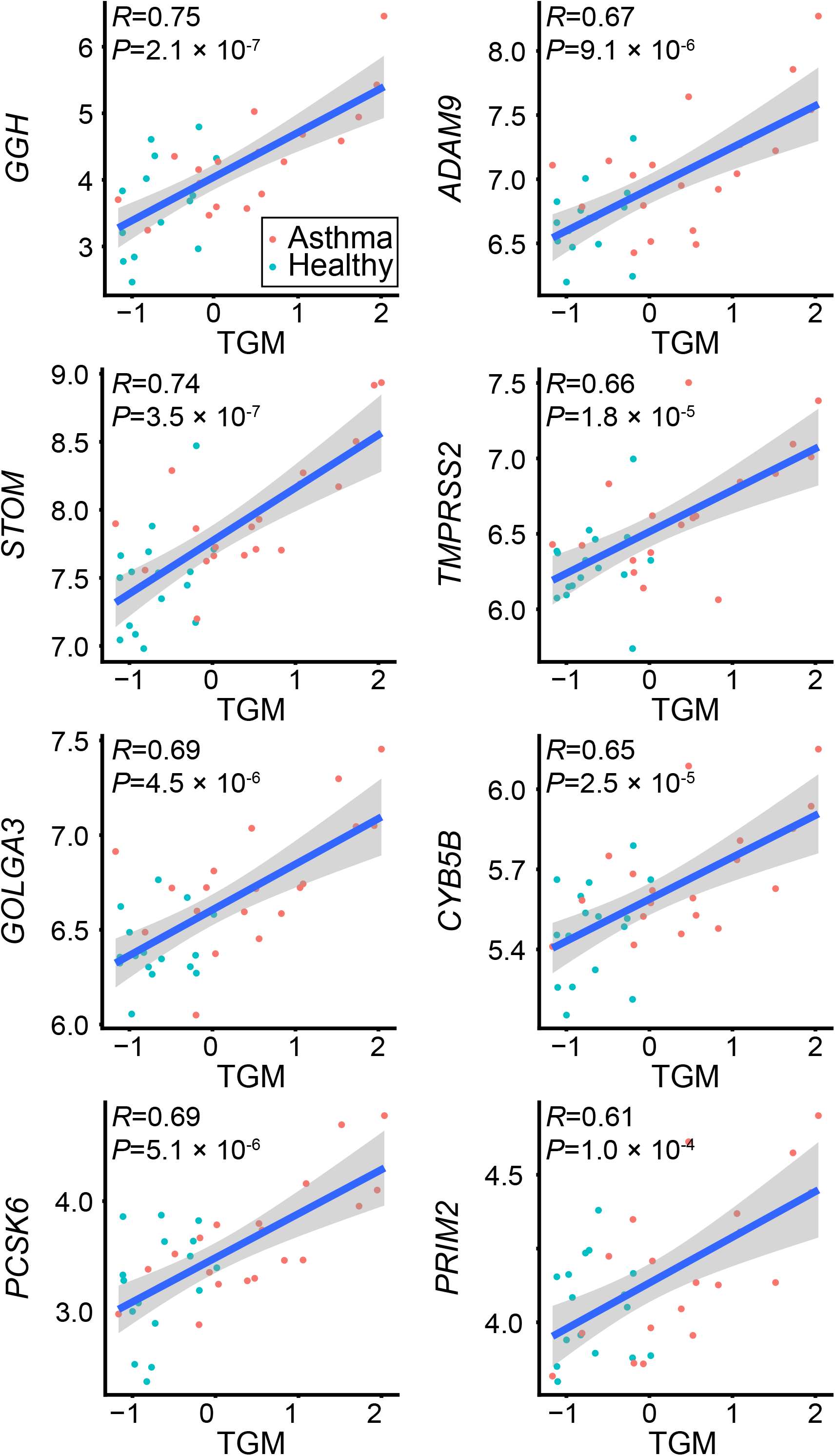
Expression of many IL-13-regulated SARS-CoV-2-associated genes correlates with an IL-13 signature in asthma. Correlation of IL-13-induced, SARS-CoV-2 associated genes with a type 2/IL-13 signature (the three gene metric, TGM) in endobronchial brushing samples from participants with asthma (red) and healthy controls (cyan). Values for gene expression represent log_2_ of normalized read counts from bulk RNA-seq. The eight SARS-CoV-2 genes with the highest Pearson’s correlations (*R*) are shown and associated *P* values are adjusted for multiple comparisons. Correlations for the full set of IL-13-induced SARS-CoV-2-associated genes are shown in Table E4.

A study of former smokers with and without COPD found that subsets of individuals with COPD also have airway epithelial gene expression changes indicative of type 2 inflammation (39). 26 SARS-CoV-2-associated genes induced by IL-13 in HBECs were measured in that study, and 21 of those positively correlated with the type 2 gene expression score developed for that study (Pearson’s *R* > 0). In 16 cases this association was statistically significant (adjusted *p* < 0.05) and remained so in a model that included age and sex (Figure E4; Table E4). Taken together, our data indicate that many IL-13-induced SARS-CoV-2-associated gene changes seen in the HBEC culture model recapitulate alterations seen in epithelial cells from individuals with asthma or COPD and type 2 inflammation.

### IL-13 protects HBECs from SARS-CoV-2 infection

We next investigated whether stimulation with IL-13 protected HBECs from SARS-CoV-2 infection by quantifying viral RNA 48 h after viral inoculation. Mucus produced by HBECs was left in place or was removed by washing the apical surface immediately prior to inoculation. In the first experiment, we found that the presence of mucus decreased the amount of SARS-CoV-2 RNA detected after infection of unstimulated cells by 74% compared with cells infected after removal of mucus (Fig. 4A). Pre-stimulation with IL-13 markedly reduced levels of SARS-CoV-2 RNA when infections were performed after removal of mucus (95% reduction) and when cells were infected in the presence of mucus (97% reduction). In a second experiment with a different HBEC donor (Fig. 4B), mucus was more effective in inhibiting infection (90-97% reduction for three different viral inocula). In cells infected after removal of mucus, IL-13 pre-stimulation reduced viral RNA by 82-92%. Since both mucus and IL-13 pre-stimulation had large effects in this donor, it was not clear whether mucus and IL-13 had additive effects in this experiment.

**Figure 4.**
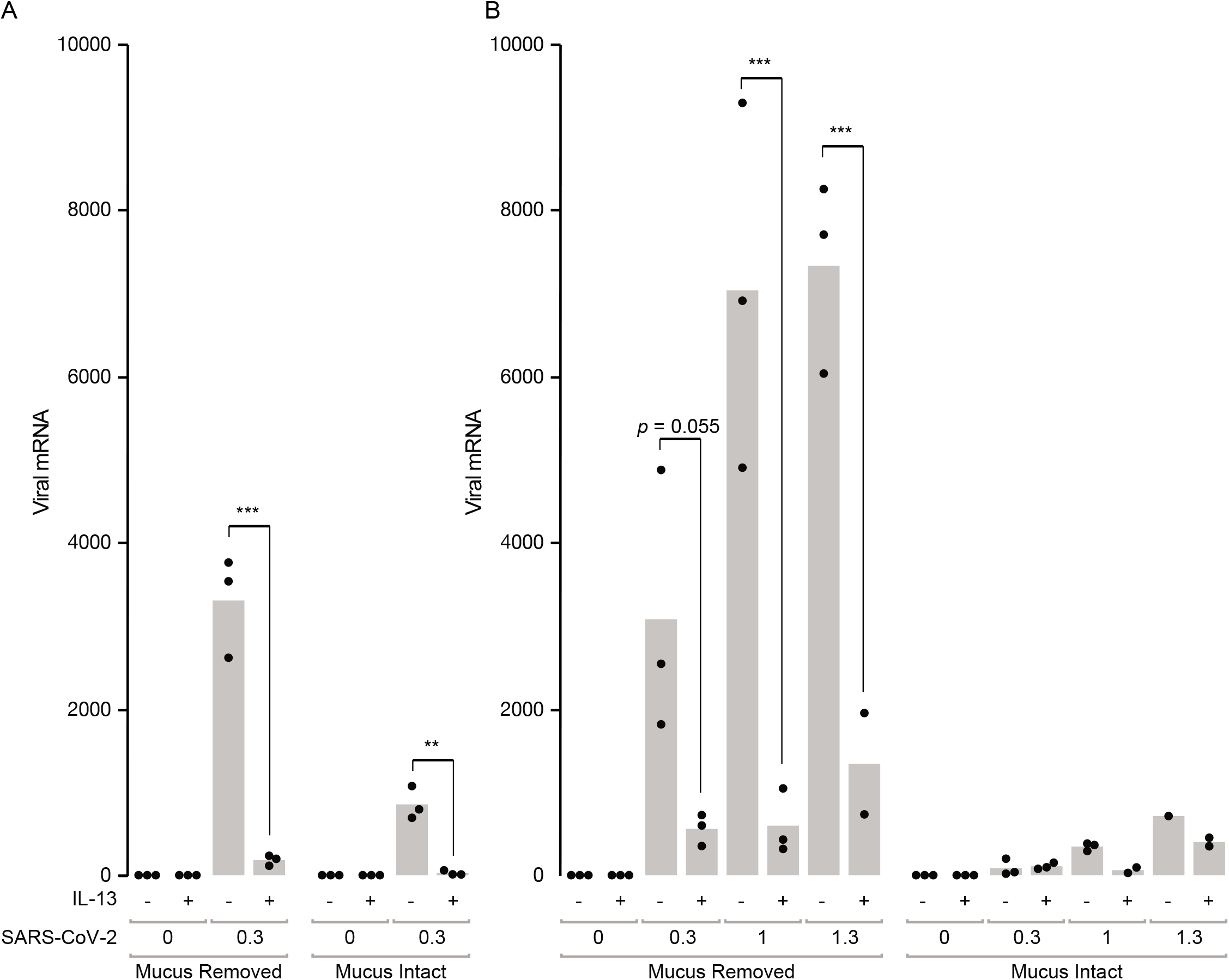
IL-13 stimulation and mucus reduce SARS-CoV-2 virus RNA levels in infected HBECs. (**A**) HBECs from a single donor were left unstimulated (–) or stimulated with IL-13 (+), washed with a DTT-containing solution (Mucus Removed) or left unwashed (Mucus Intact), and inoculated with SARS-CoV-2 (0.3 plaque forming units [pfu] based on titration in Vero E6 cells). SARS-CoV-2 mRNA was measured 48 h after infection. (**b**) In a second experiment, cells from a different donor were studied using the same protocol, except that three different inocula (0.3, 1.0, and 1.3 pfu) from another virus preparation were used. Each point represents a separate Transwell culture (n = 3 per condition except as shown). **, *p* < 0.01; *** *p* < 0.0001 for the effects of IL-13 by ANOVA with Tukey-Kramer post-tests. For cells not stimulated with IL-13, viral RNA load was lower in infections performed with mucus intact compared with infections performed with mucus removed (*p* < 0.0001 for all viral inocula in both experiments, except for *p* = 0.01 for the 0.3 pfu inoculum in the second experiment, by ANOVA with Tukey-Kramer post-tests). For viral RNA, 1 unit represents the amount of viral RNA present in 1 pfu from the viral stock, based on titration in Vero E6 cells.

We also assessed the effects of mucus and IL-13 on double-stranded RNA (dsRNA), which is produced during SARS-CoV-2 viral replication. dsRNA was not detected in uninfected cells but was detected within isolated cells or clusters of cells in unstimulated SARS-CoV-2-infected cultures. Analysis of cultures inoculated with varying amounts of virus after removal of mucus revealed a total of 27 dsRNA-stained foci with a mean volume of 6958 µm^3^ (Fig. 5). In contrast, only 3 foci (mean 1246 µm^3^) were seen in paired cultures inoculated without removing mucus (Fig. E5). In cultures pre-stimulated with IL-13, no foci were observed whether or not mucus was removed. The observation that relatively small amounts of viral RNA were detectable in IL-13-stimulated cultures (Fig. 4) but dsRNA staining was not evident under these conditions (Fig. 5 and Fig. E5) might indicate that dsRNA staining is less sensitive than qRT-PCR for viral RNA. Alternatively, it is possible that IL-13 completely prevented viral replication, and that viral RNA detected in IL-13-stimulated cells was residual RNA from the viral inoculum. Most dsRNA-containing infected cells co-stained for the ciliated cell marker acetylated-α-tubulin, although dsRNA was occasionally seen in non-ciliated cells that stained for mucins. Since PCR analysis of viral RNA indicated that the protective effects of mucus were greater in donor 2, it is noteworthy that mucin expression differed between the two donors. In the absence of IL-13, donor 1 cultures had MUC5B-containing cells but no detectable MUC5AC-containing cells (Fig. 5A and Fig. E5A), whereas donor 2 cultures had both MUC5AC- and MUC5B-containing cells (Fig. 5B and Fig. E5B). IL-13 stimulation increased MUC5AC in cells from both donors and caused an obvious decrease in MUC5B in donor 1; these effects of IL-13 are consistent with those observed in our previous studies (25) and people with type 2 asthma (19). Based on viral RNA measurements and dsRNA staining, we conclude that both the presence of a mucus gel and IL-13 stimulation reduced viral infection, and that the effects of IL-13 were seen even after removal of the mucus layer prior to infection.

**Figure 5.**
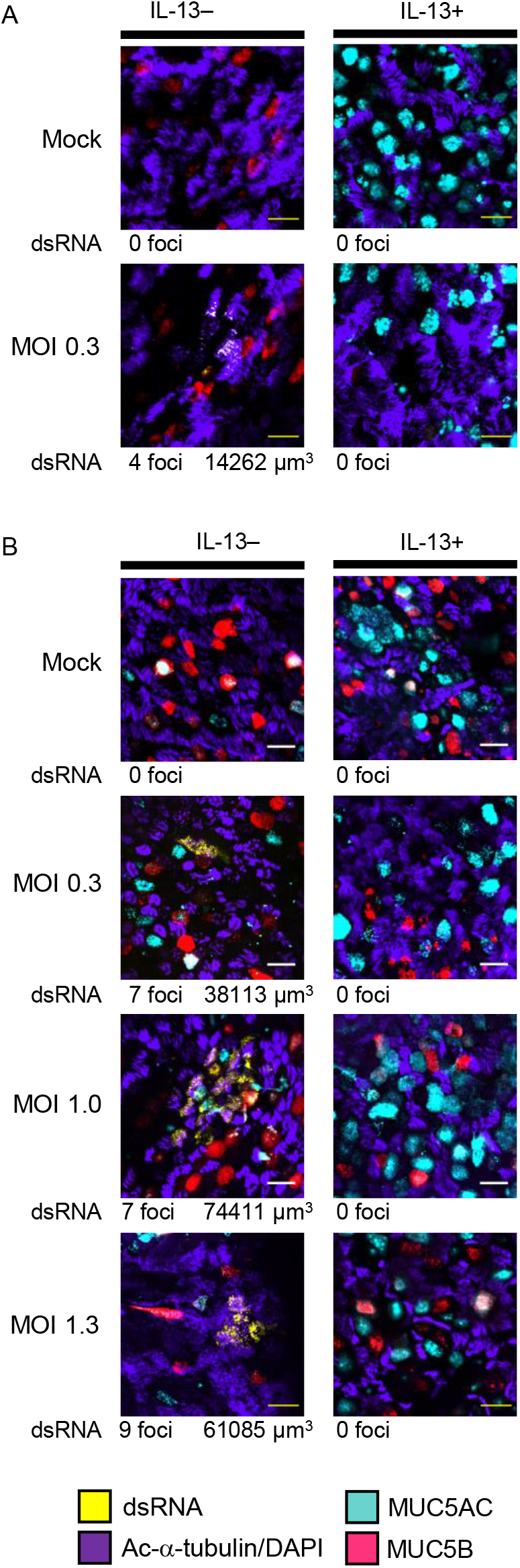
IL-13 stimulation reduces SARS-CoV-2 replication in HBECs. Additional HBEC cultures derived from cells from donor 1 **(A)** and donor 2 (**B)** were inoculated with virus after removal of mucus as part of the same experiments shown in Fig. 4. After 48 h, cells were stained with antibodies against dsRNA (yellow); the ciliated cell marker acetylated alpha tubulin (Ac-α-tubulin) and DAPI (both imaged in the same channel, purple); MUC5B (red); and MUC5AC (cyan). We surveyed the entire sample (16.6 µm^2^) for dsRNA staining and acquired stacks encompassing each dsRNA-stained focus. Numbers of dsRNA-stained foci and total volumes of dsRNA staining are shown below representative images for each condition. Scale bar = 20 µM.

## Discussion

Our studies reveal that IL-13 stimulation of HBECs affects expression of many SARS-CoV-2-associated genes and substantially inhibits SARS-CoV-2 infection of these cells. Genes encoding the large majority of SARS-CoV-2-interacting proteins identified in a previous study of HEK293T cells were expressed in HBECs. Expression of many SARS-CoV-2-associated genes differed between basal, ciliated, and secretory cells, potentially affecting how these cell types respond to SARS-CoV-2 infection. Many IL-13-induced SARS-CoV-2-associated gene expression changes we detected in culture were also seen in bronchial epithelium obtained directly from individuals with type 2 high asthma. This provides a plausible mechanism for protection against COVID-19, although the impact of asthma on COVID-19 risk is still incompletely understood and other factors may also influence COVID-19 risk in individuals with asthma (13–16). We also found significant associations of many IL-13-induced SARS-CoV-2-associated genes with type 2 inflammation in a large group of smokers with and without COPD, suggesting that the effects of IL-13 on SARS-CoV-2 risk may also be relevant in some individuals without asthma. The effects of IL-13 on SARS-CoV-2-associated genes were clearly different than the effects of IFN-α, suggesting that these two cytokines induce different antiviral mechanisms. In experiments that established the inhibitory effect of IL-13 on SARS-CoV-2 infection of epithelial cells, we found evidence that another barrier component, the mucus gel, also provides protection against infection. Taken together, these studies provide insights into airway epithelial responses that can protect against SARS-CoV-2 and might influence COVID-19 susceptibility and severity in individuals with asthma or other airway diseases.

IL-13 had a substantial effect on SARS-CoV-2 infection of HBECs as demonstrated by measurements of viral RNA and dsRNA following viral inoculation. Prior studies report a variety of effects of asthma and IL-13 on development of illnesses caused by other viruses. IL-13 can increase susceptibility of HBECs to rhinovirus infection by suppressing induction of interferons (40–42), although another study reported that prolonged pre-treatment with IL-13 of HBECs reduced rhinovirus infection (24). Mice with acute allergic airway inflammation (43) and people with pre-existing asthma (44) are reportedly protected from H1N1 influenza. Studies in IL-13-overexpressing transgenic mice and IL-13-deficient mice showed that IL-13 reduced respiratory syncytial virus replication and severity of illness (23). While effects of IL-13 on the airway epithelium are an important contributor to asthma pathogenesis, it is intriguing to speculate that IL-13 responses may have evolved at least in part to provide protection against viral infections. The finding that levels of IL-13 and the related type 2 cytokine IL-4 were higher in patients with moderate COVID-19 compared with severe COVID-19 or healthy controls is also consistent with an antiviral role for these cytokines (45).

Many mechanisms might account for IL-13-driven inhibition of SARS-CoV-2 infection. A recent study identified 65 interferon-stimulated genes that mediate restriction of SARS-CoV-2 infection (46), illustrating how a single cytokine can activate a large set of antiviral pathways. We found that gene expression changes induced by IL-13 were quite distinct from those induced by IFN-α, suggesting that these cytokines activate different antiviral pathways. We confirmed prior studies (20, 21) showing that IL-13 induced a decrease in expression of the SARS-CoV-2 receptor *ACE2*, which could contribute to decreased infection. However, the reduction in *ACE2* expression was modest compared with the effects of IL-13 on infection, suggesting that other IL-13 effects should also be considered. As in the previous reports, we found that IL-13 increased expression of *TMPRSS2*, a protease that is important for viral entry. However, we found that the IL-13 effects were cell type-dependent: *TMPRSS2* expression was increased in secretory cells but decreased in ciliated cells. Since ciliated cells were the primary cell type infected in our experiments, it is possible that decreased *TMPRSS2* in ciliated cells contributed to an overall reduction in infection. Many other host cell factors influence viral entry, RNA synthesis and translation, and egress, and further studies will be required to determine which of these contribute to the antiviral effects of IL-13.

Our studies provided clear evidence that the mucus barrier produced by HBECs in cell culture inhibits SARS-CoV-2 infection. Airway mucus is a complex hydrogel that derives its characteristic viscoelastic properties from the mucin glycoproteins MUC5B and MUC5AC (47). Prior studies establish that mucins play important roles as restriction factors for other viruses, including influenza (48, 49). We found that SARS-CoV-2 infection was decreased when mucus gels were left in place at the time of viral inoculation. We studied two subjects with distinct patterns of mucin expression and found somewhat different levels of protection from the gel.

While further studies are clearly required to investigate this further, this result suggests that differences in mucus gels are also likely to be important in SARS-CoV-2 infection. Changes in airway mucus volume, composition, and organization are prominent features of many airway diseases, including asthma (47). IL-13 is an important regulator of mucins, and we speculate that IL-13-driven increases in MUC5AC, which results in tethering of the mucus gel to the epithelium (25), might contribute to IL-13-induced inhibition of SARS-CoV-2 infection. However, this is unlikely to completely account for the antiviral effect since IL-13 inhibited viral infection even when the mucus gel was removed immediately prior to inoculation.

Our study has some important limitations. While we focused on a set of SARS-CoV-2-associated genes that have been defined in previous studies, other IL-13-regulated genes are also likely to be important for anti-viral effects. Some IL-13-regulated genes we identified in cell culture were not associated with a type 2 signature in cells from individuals with asthma or COPD, reflecting the influence of other factors, including other asthma mediators, or differences in IL-13 responses in cell culture versus *in vivo*. As individual genes that contribute to inhibition of viral infection in HBECs are identified, it will be important to specifically examine the expression of those genes in asthma and COPD. Our HBEC infection studies used only one strain of SARS-CoV-2 and cells from only two donors, and further experiments with additional strains and more donors (including donors with asthma), will be required to better understand the interactions between virus, epithelial cells, and IL-13. Finally, our infection model focuses solely on the role of epithelial cells, but the effects of IL-13 on other cell types found in the lung are also deserving of further study.

In conclusion, we found that the central asthma mediator IL-13 has a strong inhibitory effect on SARS-CoV-2 infection of HBECs. The mechanisms that account for this effect are unknown, but widespread effects of IL-13 on expression of SARS-CoV-2 associated genes that are distinct from those induced by interferons suggest that some of these mechanisms may be novel. While the use of IL-13 itself as a therapeutic may well be prevented by the pro-asthmatic effects of this cytokine, identification of IL-13-induced antiviral pathways could help address the urgent need for development of novel targeted treatments for COVID-19.

## Acknowledgements

We thank Paul Wolters (UCSF) for providing HBECs from transplant recipients, Khadija Ghias and Dingyuan Sun for assistance with HBEC culture, Melanie Ott (Gladstone Institute of Virology), Semil Choksi (UCSF), and Jeremy Reiter (UCSF) for providing advice and the staff of the UCSF Center for Advanced Light Microscopy for technical assistance.

