## Supplementary material for "The type 2 asthma mediator IL-13 inhibits SARS-CoV-2 infection of bronchial epithelium": Online Supplemental Document

**Supplementary methods**

**Human bronchial epithelial cell (HBEC) culture**

Primary HBECs from 13 individuals were cultured at air-liquid interface (ALI) as previously described (1, 2). Characteristics of the individuals are given in Table E1; samples were obtained either from donor organs not used for transplantation or from explanted organs from transplant recipients. HBECs were cultured on 12-mm Transwell inserts (Corning, Corning, NY, USA) for 3-4 weeks at ALI to obtain a well-differentiated epithelium. For cytokine stimulation, cultures were stimulated by addition of cytokines (10 ng/ml) to the basolateral medium for the final 24 h or 7 d of culture: IL-13 (Peprotech, Rocky Hill, NJ, USA), 7 d; IFN-α (R&D Systems, Inc., Minneapolis, MN, USA), 24 h; IFN-γ (R&D Systems, Inc ), 24 h; IL-17 (Peprotech), 7 d. The UCSF Committee on Human Research approved the use of HBECs for these studies.

**Bulk RNA-seq**

RNA was isolated from cytokine stimulated HBECs and bulk RNA-seq was performed as previously described (3, 4). We have previously reported other analyses based upon these data from unstimulated cells and from cells stimulated with individual cytokines (3, 4); data from cells treated with a combination of IL-13 and IFN-α have not been previously reported. All samples were cultured and analyzed as a part of a single experiment performed with cells derived from six adult donors whose lungs were not used for transplantation.

***ACE2* splicing analysis**

Differential transcript analysis was performed by first pseudo mapping reads to the Ensembl cDNA and ncRNA annotation build v103 using Kallisto [v0.46.1; (5)]. Transcript abundance estimates were then imported and corrected for transcript length using tximport (v1.14.2) and input to DESeq2 [v1.26.0; (6, 7)] to identify differentially expressed transcripts between treatment conditions. *ACE2* decoy transcripts were determined as the sum of reads mapping to MT505392.1 and ENST00000677282.

**scRNA-seq**

We used HBECs from 4 of the 6 donors used for bulk RNA-seq for scRNA-seq experiments. Cells were cultured without cytokine or with IFN-α, IL-13, IL-17, or the combination of IFN-α and IL-13. Single cell suspensions were generated by washing the apical surface briefly with PBS containing 10 mM dithiothreitol (DTT; ThermoFisher Scientific) and then incubating with 0.25% trypsin-EDTA. Cells were manually counted using a cytometer and equal numbers of cells pooled. Each pool included four samples (one from each donor, each representing a different cytokine stimulation). 10x Genomics scRNA-seq libraries were prepared (8) and sequenced using Illumina NovaSeq 6000 S4 flow cells. Resulting sequence reads were processed into a cell-by-gene counts matrix using 10x Genomics Cell Ranger software. Single nucleotide polymorphisms that were unique to one of the four donors were identified from the bulk RNA-seq data following the GATK best practices (9) and used to assign each cell to the appropriate donor using Demuxlet (10). Demultiplexed droplets uniquely assigned to a single individual were then processed using Seurat (11, 12) with the following filters: nFeature_RNA >= 500 & nFeature_RNA < 7500 & nCount_RNA > 1200 & percent.mito < .5 & percent.ribo < .35. The top 10,000 variable genes were carried forward for normalization, unsupervised clustering, UMAP visualization, and cell type identification. Normalized data was scaled while regressing out the 10x Genomics well effect, percent mitochondrial reads, percent ribosomal reads and the number of unique molecular indexes per cell. The untreated cells were then isolated for clustering and cell type identification. These untreated cells then served as the reference dataset for Seurat’s reference-based multiple dataset integration using SCTransform with label transfer in order to use anchor genes from the untreated cells to identify cell types across the other four treatments. Cell identities were then returned to the original Seurat object prior to SCTransform for further investigation. We used Seurat’s FindAllMarkers to identify cell type marker genes and the MAST statistical framework (13) to identify general and cell-type-specific cytokine effects.

**Analysis of cell type-specific expression of SARS-CoV-2-associated genes in bronchial epithelial cells**

To determine whether SARS-CoV-2-associated genes that were found to be differentially expressed in basal, secretory, and ciliated cells from HBEC cultures were also differentially expressed in freshly isolated cells from human bronchi, we examined a publicly available collection of transcriptomic profiles derived from bronchial biopsies from the airways of seven healthy donors (14). The processed scRNAseq dataset containing metadata and gene counts information were downloaded as .h5ad files from https://www.covid19cellatlas.org/. We analyzed these datasets using the same assignment of cell types the original authors used. Mean gene expression, number of cells in each cell type and fraction of cells expressing the gene were obtained using the Python module Scanpy. Further analysis and plotting was done using R v4.1.

**Asthma and COPD gene expression analyses**

For analyses of gene expression in asthma, we used gene expression data from the Mechanisms of ASThma Study (MAST) (3). Participants with asthma (n = 19) were required to have a positive methacholine bronchoprovocation test, and healthy controls (n = 16) were required to have no history of asthma or allergies. Subjects could not have used steroids in the 6 weeks prior to enrollment. Subjects underwent research bronchoscopy and endobronchial brushings were analyzed using bulk RNA-seq. We correlated the three gene mean (TGM) in the both clinical datasets with SARS-CoV-2-associated genes differentially expressed following IL-13-stimulation using Pearson’s correlation coefficient. To determine whether associations with the TGM remain after adjustment for age and sex, we used the linear regression model TGM ~ gene expression + age + sex.

For analyses of gene expression in COPD, we used data from a previous study of 238 current and former cigarette smokers with and without COPD (15). A small subset of 17 subjects reported a history of asthma. Subjects underwent research bronchoscopy and endobronchial brushings were analyzed using DNA microarrays. Analyses of correlations with IL-13-regulated SARS-CoV-2-associated genes were performed as in the MAST analysis, except that we used a type 2 score (T2S) previously developed for use with this dataset (16) rather than the TGM as the index of type 2/IL-13 activity.

**SARS-CoV-2 propagation and titration**

SARS-CoV-2 virus (USA-WA1/2020 strain) and Vero E6 cells were provided by Dr. Melanie Ott (Gladstone Institute of Virology). For virus propagation, Vero E6 cells were cultured in Dulbecco's Modified Eagle Medium (UCSF Media Production) supplemented with 10% fetal bovine serum (Corning), penicillin/streptomycin (UCSF Media Production), and L-glutamine (Corning) in a 37° C, 5% CO_2_ incubator. Vero E6 cells were infected with SARS-CoV-2 virus, incubated supernatant was collected at 72 h. The virus was aliquoted and stored at -80° C. Viral titer was quantified using a plaque assay in Vero E6 cells. 10-fold dilutions of the virus stock were added to Vero E6 cells in a 12-well plate for 1 h, after which an overlay of 1.25% Avicel RC-591 in DMEM was added. The cells were incubated at 37° C, 5% CO_2_ for 72 h. The cells were fixed with 10% formalin, stained with crystal violet, and washed with water. The plaques were counted to determine the titer of the virus stock. All work was done under BSL3 conditions.

**Quantitative real-time PCR (qRT-PCR)**

48 h after SARS-CoV-2 infection, RNA was isolated from HBECs with the Zymo RNAEasy Micro kit. Viral RNA was reverse transcribed using SuperScript III First-Strand Synthesis System (ThermoFisher Scientific). cDNA was analyzed by qRT-PCR using PowerUp SYBR Green (ThermoFisher Scientific) and primers specific for SARS-CoV-2 Orf1b [HKU-ORF1b-nsp14_F: TGGGGYTTTACRGGTAACCT, HKU-ORF1b-nsp14_R: AACRCGCTTAACAAAGCACTC (17)] and detected using 7900HT Fast Real-Time PCR System (ThermoFisher Scientific). Serial dilutions of RNA isolated from a viral stock were used to generate a standard curve. Statistical analysis was performed using ANOVA with Tukey-Kramer post-test in JMP (SAS Institute, Cary, NC).

**Immunofluorescence**

48 h after SARS-CoV-2 infection, HBECs were fixed in 4% paraformaldehyde [PFA; (ThermoFisher Scientific)] for 30 min, washed twice and then stored in PBS at 4° C until stained. Filters were removed from inserts and then simultaneously blocked and permeabilized using 5% normal goat serum (Jackson ImmunoResearch Laboratories, Inc, West Grove, PA, USA) and 0.1% Triton X100 in PBS for 30 min at room temperature. Filters were incubated with primary antibodies against acetylated-α-tubulin (1:150; Santa Cruz Biotechnology, Inc,, Dallas, TX, USA), MUC5B (1:150; Santa Cruz), MUC5AC (1:200; ThermoFisher Scientific) and dsRNA (Jena Bioscience, Thuringia, Germany) overnight at 4° C. Filters were than 4,6-diamidinio-2-phenylindole (DAPI) was used to counterstain nuclei (1:1000). washed in TBST and incubated with goat secondary antibodies (1:200; Jackson ImmunoResearch Laboratories, Inc) for 2-3 h at room temperature; as acetylated-α-tubulin is located apically and nuclei basally, goat anti-mIgG2b-BV421 was used to detect acetylated-α-tubulin in the same channel as the DAPI staining, despite their overlapping spectra. Filters were washed again and mounted in Fluoromount G (SouthernBiotech, Birmingham, AL, USA). 4,6-diamidinio-2-phenylindole (DAPI) was used to counterstain nuclei (1:1000). Slides were imaged using a Yokagawa CSU22 spinning-disk confocal microscope connected to a Nikon Ti-E and a dry 40x objective at the Center for Advanced Light MIcroscopy, UCSF. We surveyed the entire sample (16.6 µm^2^) for dsRNA staining and acquired stacks comprising 0.6 µM slices spanning the thickness of the epithelium at each focus of dsRNA staining, and at random locations for inserts lacking any dsRNA staining.

**Image analysis**

Images were analyzed using ImageJ2 (18). Nd2 files were read using Bioformats. Stacks were split into individual channels and thresholded by the Otsu method (19). For composite images, channels were recombined and converted to rgb images. The volume of dsRNA staining was determined from the thresholded dsRNA channel using measurements of the dsRNA-stained areas in all slices from each stack.

**Supplementary Tables**

**Table E1: HBEC donor characteristics**

| Donor | Origin | Sex | Age | Race | Hispanic | Disease | Smoking Status | Bulk RNA-seq | scRNA-seq | Viral infection |
| --- | --- | --- | --- | --- | --- | --- | --- | --- | --- | --- |
| 12-43 | NA | Male | 42 | White | Y | No airway disease | Non-smoker | + | + | - |
| 13-26 | TB | Male | 40 | White | N | No airway disease | Non-smoker | + | + | - |
| 13-28 | TB | Male | 51 | White | Y | No airway disease | Non-smoker | + | + | - |
| 13-33 | TB | Male | 50 | White | N | Asthma | Smoker | + | + | - |
| 13-23 | TB | Male | 59 | White | N | No airway disease | Non-smoker | + | - | - |
| 13-24 | B | Female | 67 | White | Y | No airway disease | Non-smoker | + | - | - |
| 11-35 | B | Male | 49 | White | N | ILD | NA | - | - | - |
| 13-27 | TB | Male | 46 | White | N | No airway disease | Smoker | - | - | - |
| 13-43 | NA | Male | 14 | NA | NA | ILD | NA | - | - | - |
| 14-11 | B | Female | 47 | White | N | ILD | NA | - | - | - |
| 18-36 | B | Female | 56 | White | N | ILD | NA | - | - | - |
| 10-75 | B | Male | 38 | NA | N | ILD | Non-smoker | - | - | + |
| 14-30 | B | Male | 64 | White | N | ILD | Smoker | - | - | + |

* NA, not available; TB, tracheobronchial; B, bronchial; Y, yes; N, No; ILD, Interstitial Lung Disease

*Tables E2-E4 are provided as Excel files.*

**Table E2: Bulk RNA-seq data for SARS-CoV-2-associated gene expression in unstimulated and cytokine-stimulated HBECs**

**Table E3: scRNA-seq data for SARS-CoV-2-associated gene expression in unstimulated and cytokine-stimulated HBECs**

**Table E4: Associations between IL-13 signatures and expression of IL-13-regulated SARS-CoV-2-associated genes in asthma and COPD**

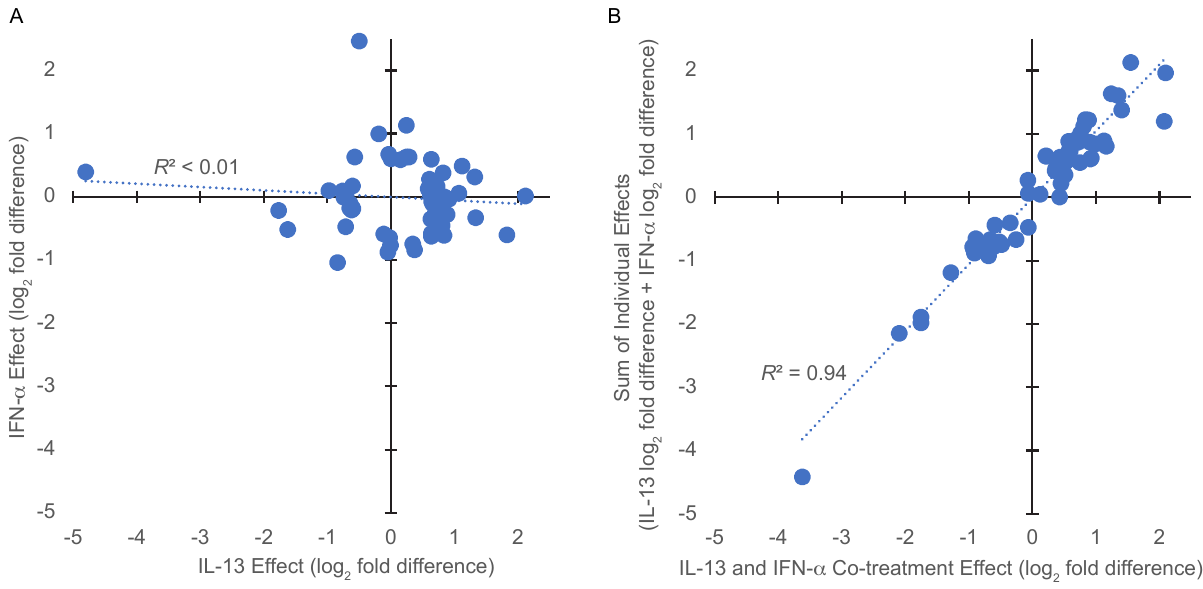

**Figure E1. Comparison of IL-13 and IFN-α effects on expression of SARS-CoV-2-associated genes.** Figures show effects for all SARS-CoV-2-associated genes that were affected by either IL-13 or IFN-α stimulation (absolute fold difference > 1.5, FDR *q* < 0.05). (**A**) Comparison of IL-13 and IFN-α effects. (**B**) Comparison of the effects of co-treatment with IL-13 and IFN-α with the sum of the effects of treatment with each individual cytokine. *R,* Pearson’s correlation coefficient.

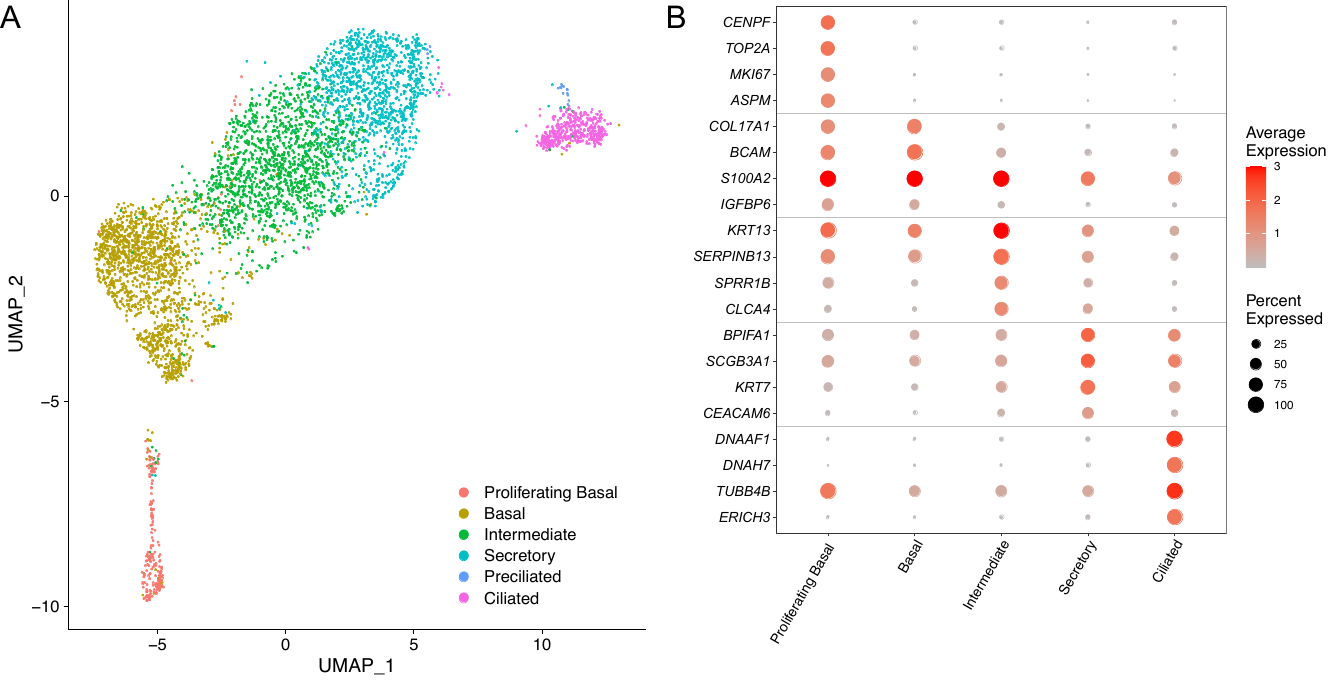

**Figure E2. Identification of cell types from scRNA-seq data. (A)** Uniform Manifold Approximation and Projection (UMAP) plot showing cell clustering and cell type assignments. **(B)** Expression of marker genes in the five major subsets.

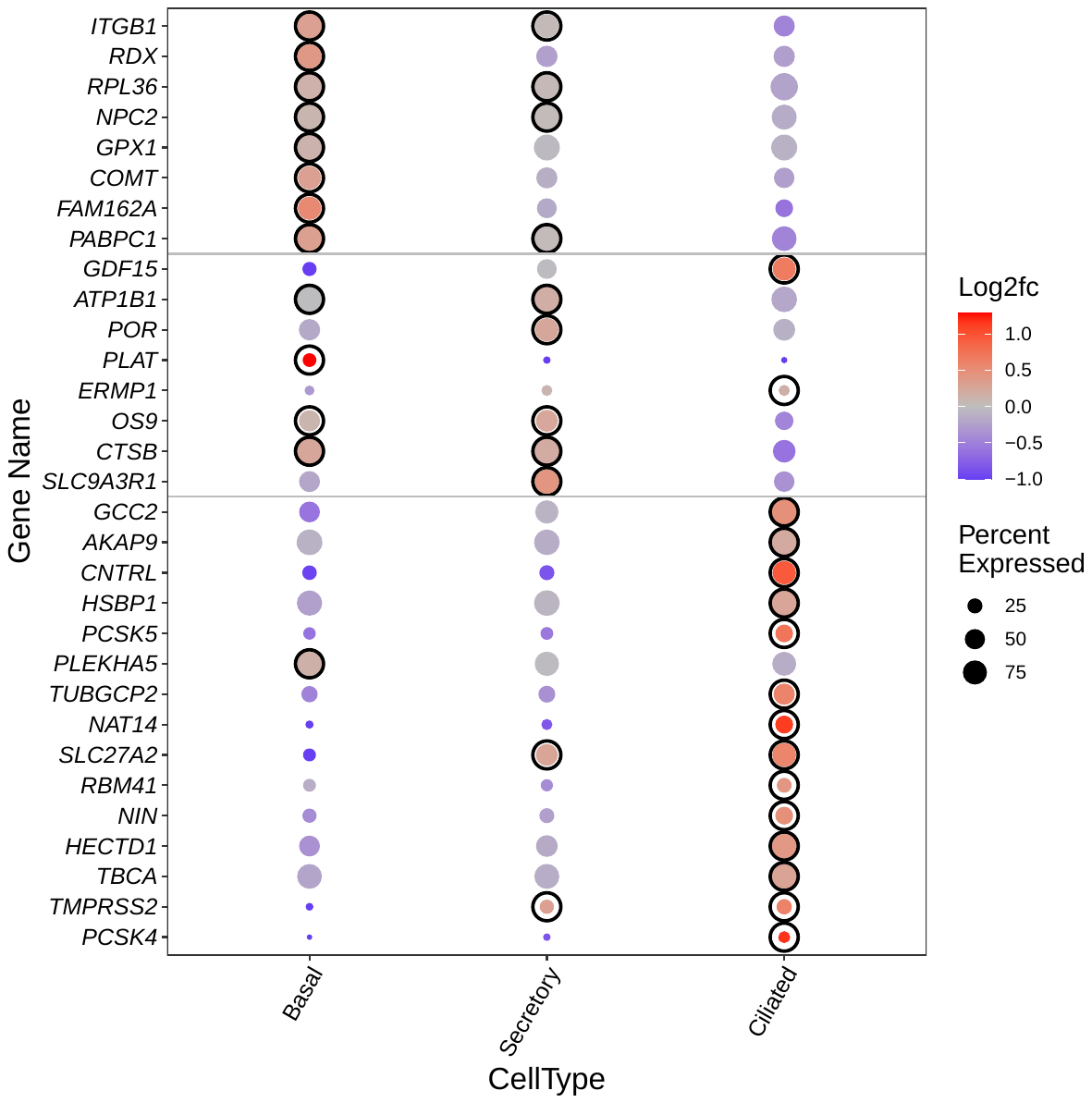

**Figure E3. Cell type-specific expression of SARS-CoV-2-associated genes in freshly isolated cells from human bronchi.** Genes were selected from those identified as cell type-specific in the analysis of cultured HBECs. scRNA-seq data are from reference (14).

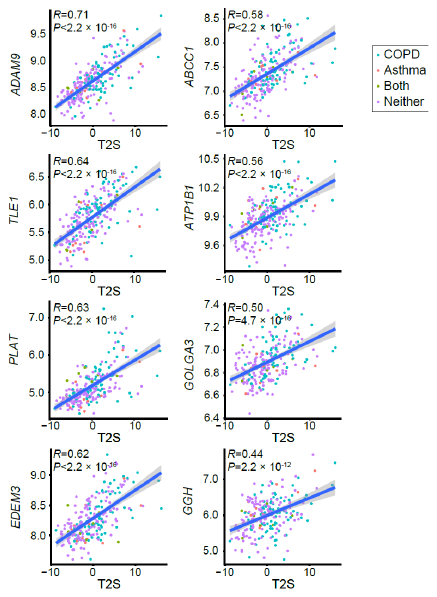

**Figure E4. Expression of many IL-13-regulated SARS-CoV-2-associated genes correlates with an IL-13 signature in chronic obstructive pulmonary disease (COPD).** Correlation of IL-13-induced, SARS-CoV-2 associated genes with a type 2/IL-13 signature (type 2 score, T2S) in endobronchial brushing samples from participants with asthma (red), COPD (blue), both asthma and COPD (green) or neither (purple). Values for gene expression represent log2 of normalized read counts from bulk RNA-seq. The eight SARS-CoV-2 genes with the highest Pearson’s correlations (*R*) are shown and associated *P* values are adjusted for multiple comparisons. Correlations for the full set of IL-13-induced SARS-CoV-2-associated genes are shown in Table E4.

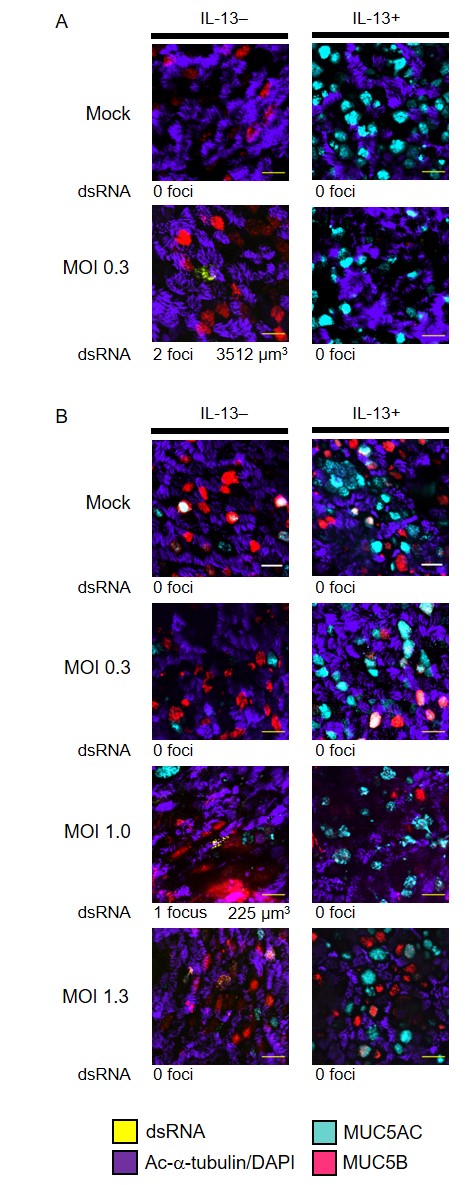

**Figure E5. Low levels of SARS-CoV-2 replication when mucus is not removed prior to inoculation HBECs.** Additional HBEC cultures derived from cells from donor 1 **(A)** and donor 2 (**B)** were inoculated with virus without removal of mucus as part of the same experiments shown in Fig. 4. After 48 h, cells were stained with antibodies against dsRNA (yellow); the ciliated cell marker acetylated alpha tubulin (Ac-α-tubulin) and DAPI (both imaged in the same channel, purple); MUC5B (red); and MUC5AC (cyan). We surveyed the entire sample (16.6 µm^2^) for dsRNA staining and acquired stacks encompassing each dsRNA-stained focus. The numbers of dsRNA-stained foci and the total volumes of dsRNA staining are shown below representative images for each condition. See Fig. 5 for comparison of cells inoculated after removal of mucus. Scale bar = 20 µM.
